# Distinct Social Behavior and Inter-Brain Connectivity in Dyads with autistic individuals

**DOI:** 10.1101/2023.05.01.538921

**Authors:** Quentin Moreau, Florence Brun, Anaël Ayrolles, Jacqueline Nadel, Guillaume Dumas

## Abstract

Autism Spectrum Disorder (ASD) is defined by distinctive socio-cognitive behaviors that deviate from typical patterns. Notably, social imitation skills appear to be particularly impacted, manifesting early on in development. This paper compared the behavior and inter-brain dynamics of dyads made up of two typically developing (TD) participants with mixed dyads made up of ASD and TD participants during social imitation tasks. By combining kinematics and EEG-hyperscanning, we show that individuals with ASD exhibited a preference for the follower rather than the lead role in imitating scenarios. Moreover, the study revealed inter-brain synchrony differences, with low-alpha inter-brain synchrony differentiating control and mixed dyads. The study’s findings suggest the importance of studying interpersonal phenomena in dynamic and ecological settings and using hyperscanning methods to capture inter-brain dynamics during actual social interactions.

## Introduction

Autism spectrum disorder (ASD) is a neurodevelopmental condition characterized by atypical social behaviors, ranging from non-verbal interactions to sophisticated social cognition (American Psychiatric Association, 2013; Grzadzinski et al., 2013; Lord et al., 2020). For instance, imitation skills in children with ASD are notably diminished (Ingersoll, 2008) while in typical development (TD), social imitation allows children to learn from others (Heyes, 2011; Ray and Heyes, 2011) but also reflects their search for belongingness (Over and Carpenter, 2013; Over, 2016).

The nature of social imitation differences in ASD remains unclear. Some studies argue for a dysfunction in the “mirror” system (Oberman et al., 2005; Yang and Hofmann, 2016), with specific structural alterations in the angular gyrus (Mengotti et al., 2013). However, recent evidence points against this hypothesis (Dumas et al., 2014; Hobson and Bishop, 2016; Heyes and Catmur, 2022). Although the brain network involved in motor imitation might be under-activated in ASD, imitation abilities do not differ from the typically developed controls (Wadsworth et al., 2017), with even evidence of imitation learning in low-function autistic children (Nadel et al., 2011). On the other hand, others argue that social imitation deficits are due to lower-level atypical social perception such as reduced sensitivity to biological motion in ASD (Mason et al., 2021). However, most findings in imitation research emerged from single participants’ experiments examining brain responses to pictures or video clips passively shown. Moreover, several problems usually labeled as general impairment of imitation in ASD, such as a narrow motor repertoire, are likely due to restricted interests and altered attention to others’ behaviors (Nadel, 2014a). Indeed, when put in a dyadic context, children with ASD show spontaneous imitation capacities and recognize when they are being imitated (Escalona, et al., 2002).

The last two decades have witnessed a “second-person” shift in social neuroscience, with the development of interactive set-ups involving more than one participant at a time (Schilbach et al., 2013). In parallel, hyperscanning (Montague et al., 2002; Cui et al., 2012) now allows the simultaneous recording of two or more individuals’ brain activities, especially while participants interact with each other (Babiloni and Astolfi, 2014). Thanks to this method, inter-cerebral correlation (IBC) and inter-brain synchrony (IBS) have been observed in various social contexts such as mutual gaze (Leong et al., 2017), shared attention (Hirsch et al., 2017), face-to-face deception (Zhang et al., 2017), social connectedness among interacting partners (Kinreich et al., 2017), empathy (Mengotti et al., 2013), verbal interactions (Hirsch et al., 2018), but also coordination (Zamm et al., 2018), interpersonal synchronization (Cui et al., 2012) and collaboration (Matusz et al., 2019). Those inter-brains communications highlight how we can no longer think of the brain as an isolated object (Bottema-Beutel et al., 2019). Consequently, to understand social imitation in ASD, we need to study individuals in bilateral and spontaneous interactions (Nadel & Pezé, 1993).

To test our hypotheses, we asked dyads to perform a social imitation task with hand movements, while we recorded their movements and hyperscanning-EEG. We contrast the results between ASD-TD and TD-TD pairs of participants, with the main hypothesis that ASD-TD dyads would show distinct behavioral and inter-brain dynamics patterns compared to TD-TD ones.

## Methods

### Participants

Forty participants, ten high-functioning adults with Autism Spectrum Disorder (7 males, 3 females; M age ± SD 33.9 ± 6.2 years; range 21–41 years) and thirty typical adults (14 males, 16 females; M age SD 28.7 ± 5.2 years; range 20–39 years), participated in the study, resulting in ten dyads in the Mixed Dyads group (ASD-TD) and ten in the Control Dyads group (TD-TD). Due to technical problems in the recordings of two dyads, the final sample consists of nine dyads in the Mixed Dyads group and nine in the Control Dyads group.

The exclusion criteria were associated with past or present neuropsychiatric and neurological disorders. All participants had normal or corrected-to-normal vision. They were right-handed (except for one individual in the ASD group). All were volunteers and gave their written informed consent according to the Declaration of Helsinki. The institutional ethical review board for Biomedical Research of the Hospital of Pitié-Salpétrière approved the experimental protocol (agreement #104-10).

The diagnosis of high-functioning ASD was established by psychiatrists and neuro-psychologists with the DSM-IV-R (American Psychiatric Association, 2002), the Autism Diagnostic Interview-Revised (ADI-R; Lord et al., 1994), the Autism Diagnostic Observation Schedule-Generic (ADOS-G; Lord et al., 2000) module 4 (mean Social-communication score = 10.8, SD = 5.77), and expert clinical evaluation. No ASD participant underwent any drug and/or intervention program or participated in another experiment during the study.

Four ASD participants studied at the university with at least three years of training, and six practiced high-level professions (graphic teacher, archivist, librarian, psychotherapist, engineer, and computer programmer). All TD participants studied at the university with at least three years of training. Academic achievement is therefore comparable between groups.

### Procedure

Each participant of a dyad sat in a separate room. They faced a 21-inch TV screen, with forearms resting on a table to prevent arms and neck movements (Figure 1A). In each experimental room, a digital video camera filmed participants’ hand gestures. These films were transmitted live to the TV screen of the other room and to the experimenter’s recording room. Thus, each participant could see the other’s hand gestures in real-time and the experimenter could control that participants followed the requested instructions. The session start was signaled by a LED light manually controlled by the experimenter via a switch.

**Figure 1.**
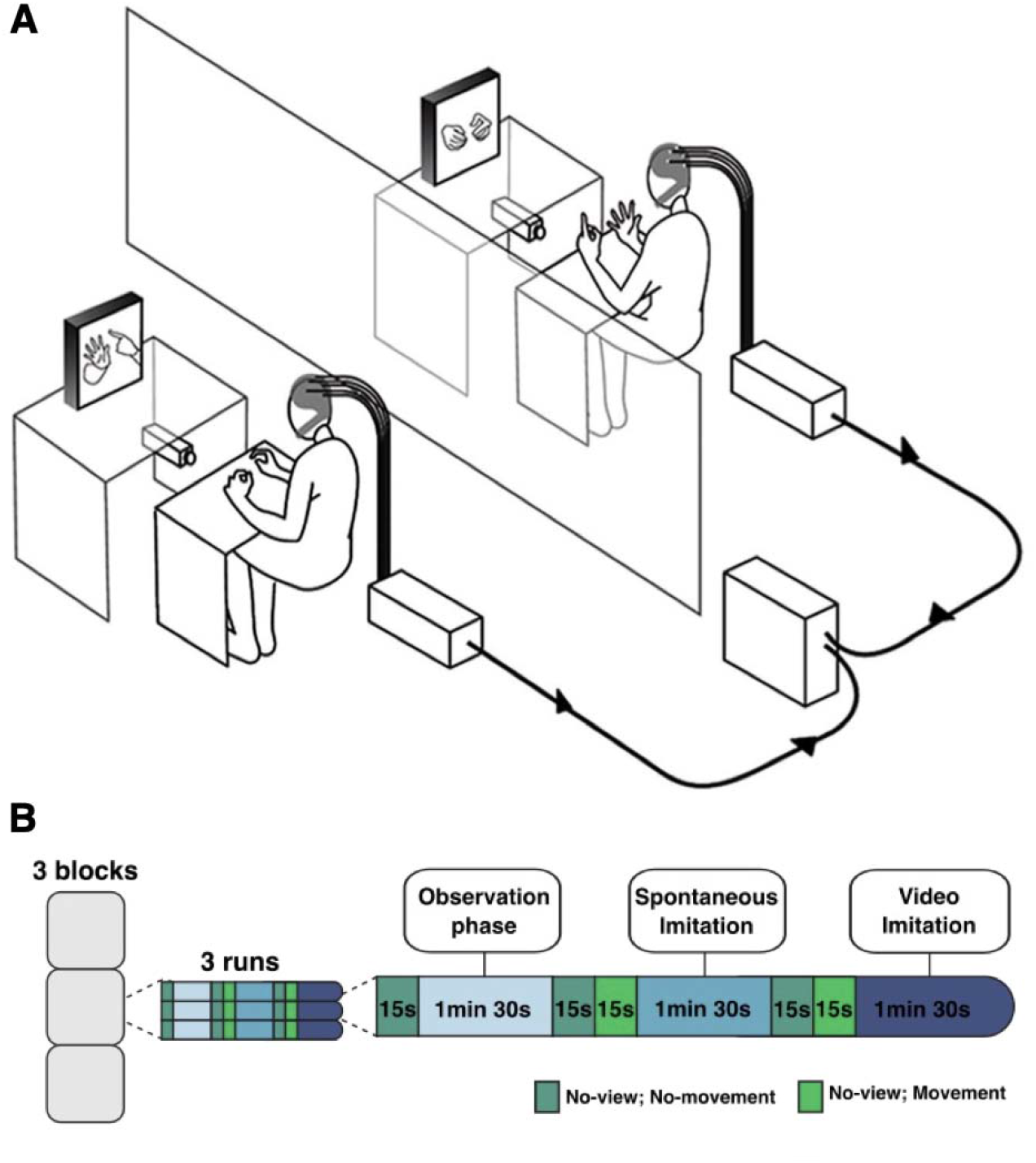
**A**. Experimental setting of the double video system and dual-EEG recording. **B**. Example of an experimental run, where an Observation phase of a prerecorded library of 20 meaningless hand gestures (1’30), a Spontaneous Imitation phase (1’30) where the participants could either produce hand gestures of their own or imitate hand gestures from the other participant transmitted by the video camera, and a Video Imitation phase (1’30) where the participants were asked to imitate a prerecorded video. Each run started with a 15s resting-state period with No-View No-Movement (NVNM), repeated between each condition. The Spontaneous and Video Imitation phases also had a period where participants were asked to produce meaningless hand gestures with no visual feedback from the other participant (i.e. No-View Movement, NVM).

The experimental protocol was divided into three blocks separated by a 10 min pause (Figure 1B). Each block comprised three runs, where a run of 3 conditions: an *Observation* phase of a prerecorded library of 20 meaningless hand gestures (1’30), a *Spontaneous Imitation* phase (1’30) where the participants could either produce hand gestures of their own or imitate hand gestures from the other participant transmitted by the video camera, and a *Video Imitation* phase (1’30) where the participants were asked to imitate a prerecorded video. Each run started with a 15s resting-state period with No-View No-Movement (NVNM), repeated between each condition. The Spontaneous and Video Imitation phases also had a period where participants were asked to produce meaningless hand gestures with no visual feedback from the other participant (i.e. No-View Movement, NVM). At the end, a short block of calibration comprised periods of blinks, jaws contraction, and head movements of 30 seconds each. All conditions were presented in a fixed order for group comparison. For further information about the design, please look at previous papers (Dumas et al., 2010, 2014).

### Hyperscanning-EEG acquisition

The neural activities of the two participants were simultaneously recorded with a dual-EEG recording system. It was composed of two Acticap helmets (Brain Products, Germany) with 64 active electrodes arranged according to the international 10/20 system. The helmets were aligned to nasion, inion, and left and right pre-auricular points. A three-dimensional Polhemus digitizer (Polhemus Inc., Colchester, VT, USA) was used to record the position of all electrodes and fiducial landmarks (nasion and pre-auricular points). The ground electrode was placed on the right shoulder of the participants and the reference was fixed on the nasion. The impedances were maintained below 10 kΩ. Data acquisition was performed using two 64-channel Brainamp MR amplifiers (Brain Products, Germany). Signals were analog filtered between 0.16 Hz and 250 Hz, amplified, and digitized at 500 Hz with a 16-bit vertical resolution in the range of ± 3.2 mV.

### Data analysis

#### Behavioral data analyses

We analyzed the video recordings of hand movements during Spontaneous Imitation and Video Imitation to define periods of time during which participants were really imitating, in contrast to non-imitative periods (based on the morphology and direction of the hand movement, see previous work for more details (Delaherche et al., 2014)). Through homemade Matlab codes, we extracted several variables. We first computed the total duration of imitative periods (*Overall Imitation*, see Figure 2A) to assess whether the task instructions were followed (i.e., how long each dyad correctly performed the task by imitating each other). Additionally, we analyzed the video recordings of hand movements during *Spontaneous Imitation*: we extracted measures of interactional synchrony, measured when the hands of the two participants started and ended a movement simultaneously, thus showing a coordinated rhythm (*Synchrony*, Figure 2B). We also distinguished *Role Symmetry*, to explore whether dyads had a balanced repartition of roles (Figure 2C), using the formula:

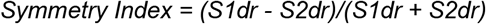

where S1dr and S2dr represent the time spent as a model by subject 1 and subject 2 respectively, b□=□0 indicating a perfect symmetry of the two roles (Dumas *et al*., *2010*).

**Figure 2.**
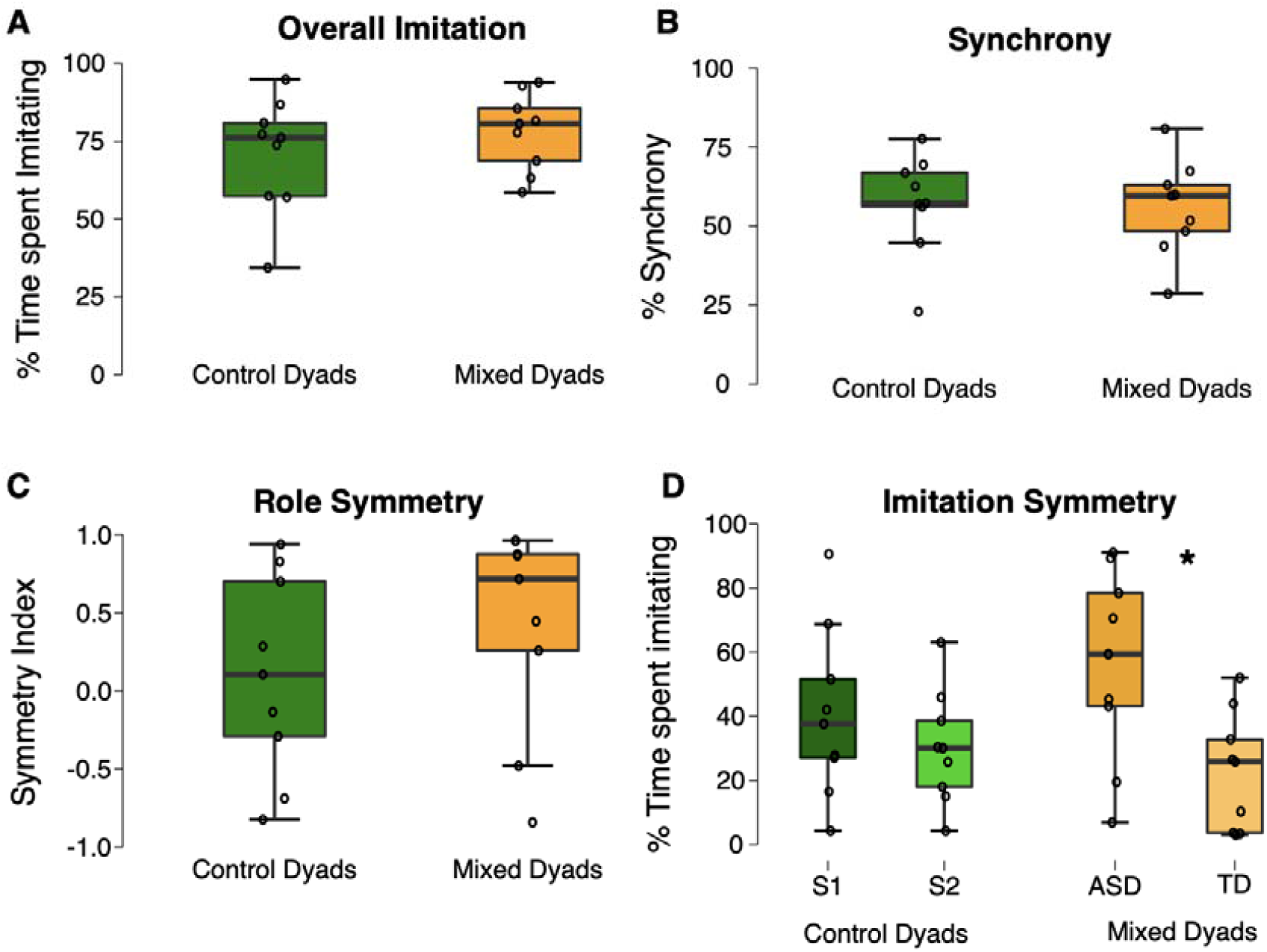
Behavioral results comparing Control and Mixed Dyads for **A**. Overall Imitation; **B**. Synchrony; **C**. Role Symmetry and **D**. Imitation Symmetry variables - * shows significant differences (*p* < 0.05).

Finally, we broke down the *Symmetry Index* values between each member of the dyad to better highlight the driving and following roles between the participants across time (*Imitation Symmetry*, reflecting the time spent imitating, see Figure 2D) within each dyad.

Due to the violation of two-sample t-test assumptions, *Overall Imitation, Synchrony*, and *Role Symmetry* variables were analyzed using non-parametric Mann-Whitney tests between Control Dyads (TD-TD) and Mixed Dyads (ASD-TD). *Imitation Symmetry* was analyzed using a mixed repeated-measure ANOVA with Group (Control, Mixed) as a between-subjects factor, and Subjects (S1/ASD, S2/TD) as a within-subjects factor.

#### EEG-Hyperscanning data preprocessing

The EEG data were initially pre-processed in Matlab, where blink, muscles, and head movement artifacts were filtered by optimal projection (FOP) methodology (Boudet et al., 2007). Visual control allowed to reject the remaining artifacts (<0.1% of the data, no difference between the two groups), and noisy EEG channels that were marked as bad (for ASD subjects: average = 1.80 ± 2.82, min = 0, max = 10; for TD subjects: average□= 1.06 ± 1.19, min□= 0, max□= 6). We then converted the EEG data from Matlab to Python and used the MNE-Python library (Gramfort et al., 2013) for further analyses and statistics on the EEG data. We also used the open-source library Hyperscanning Python Pipeline (HyPyP), based on MNE-Python, that our team implemented (https://github.com/GHFC/HyPyP, Ayrolles et al., 2021) to analyze inter-brain dynamics.

For each phase of a run (Resting-States, *Observation Phase, Spontaneous Imitation*, or *Video Imitation)*, we converted the hyperscanning-EEG data to the MNE-Python Raw data format. Then, we low-pass filtered Raw data at 2 Hz with a finite impulse response filter and created 1s epochs around fixed events. The epochs were concatenated across the blocks for each condition (*Observation Phase, Spontaneous Imitation, Video Imitation*). Epochs were cleaned for each dyad using the preprocessing HyPyP functions adapted from Autoreject (Jas et al., 2017). The process involved the rejection of all epochs marked bad for at least one participant, the rejection or interpolation of partially bad sensors per participant, and the removal of the irreparable bad sensors across participants. Thus, only sensors and epochs that were deemed “good” for the two participants were preserved.

#### Neurodynamical analyses

We defined 4 frequency-bands-of-interest: Theta [4-8], Alpha Low [8-10], Alpha High [11-13], Beta [14-31] Hz. For each frequency-bands-of-interest, we estimated the analytic signal by a multitaper and calculated the circular correlation coefficient between all inter-brain sensor pairs of a dyad. Circular Correlation (CCorr) measures the covariance of phase variance between two data streams and is more robust to coincidental synchrony (Burgess, 2013) compared to phase-locking value or phase-locking index. CCorr has seen increasing popularity and has been successfully implemented in studies investigating touch (Goldstein et al., 2018), learning (Davidesco et al., 2019), and language (Perez Repetto et al., 2017). We averaged circular correlation coefficient values across epochs and applied the Log ratio normalization mentioned above.

#### EEG-behavior preprocessing

In order to match the EEG along *Imitation Symmetry* behavioral values (i.e., distinguishing leaders and followers within the dyad), for the *Spontaneous Imitation* condition, we cropped the filtered Raw data corresponding to the task to differentiate periods of time during which: a) participant 1 was driving and participant 2 was following the hand movement b) participant 2 was driving and participant 1 was following the hand movement c) they do not really imitate each other. We epoched the raw data for each period of time with different event identities and concatenated all of them across the blocks. Concatenated Epochs were cleaned for each dyad as described above. Then, we were able to split cleaned Epochs between the periods of time we mentioned thanks to event identity. Cleaned Epochs corresponding to periods of time 1/ and 2/ were used for further analyses, and realigned alongside the same axis so that all leader and follower epochs were ordered alongside the same dimension (i.e., instead of divided by participant 1 and participant 2). We show the results of these analyses in Figure 5.

#### Statistics

We used a cluster-level statistical permutation test to contrast CCorr values between dyads and conditions. When comparing within dyads, the statistical test and threshold used in the cluster-level statistical permutation test were provided by dependent t-test (p-value = 0.025) and when comparing dyads, we used cluster-level statistics provided by one-way repeated measure ANOVA (p-value 0.025). The cluster-level statistical permutation test reduces family-wise error due to multiple comparisons by clustering neighboring quantities that exhibit the same effect. The neighborhood is corrected by space (adjacent sensors over the scalp) and frequencies (adjacent frequency bins). We assumed no sensors’ connectivity between the brains of the two participants in each dyad - however, we took into account intra-participant neighboring (inter-brain sensors’ pairs including common or neighboring sensors in one of the two participants). The sum of t values in a given cluster was used for cluster-statistic. Clusters’ p-value was estimated through the distribution of cluster-statistics from randomizations of the dataset (Maris *et al*., 2007; Gramfort *et al*., 2013) 5000 permutations, p-value set at 0.025).

## Results

### Behavioral measures

#### Overall Imitation

The Mann-Whitney test revealed no significant difference between Control Dyads (M = 70.959, SD = 18.405) and Mixed Dyads (M = 78.111, SD = 12.386) for Overall Imitation along trials (W=30.00, *p =* 0.387, rrb = 0.259, see Figure 2A).

#### Synchrony

The Mann-Whitney test revealed no significant difference between Control Dyads (M = 57.173, SD = 15.895) and Mixed Dyads (M = 55.913, SD = 15.022) for Synchrony measure (W=43.00, *p =* 0.863, rrb = 0.062, see Figure 2B).

#### Role Symmetry

The Mann-Whitney test revealed no significant difference between Control Dyads (M = 0.102, SD = 0.644) and Mixed Dyads (M = 0.419, SD = 0.663) for the Symmetry Index (W=27.00, *p =* 0.258, rrb = 0.333, see Figure 2C).

#### Imitation Symmetry

Distinguishing between TD in the Controls Dyads and TD and ASD members within Mixed Dyads, we see a significant difference in time spent imitating (*F*(3, 32) = 3.387, *p =* 0.030, η_p_^2^ = 0.241). Post-hoc tests only revealed a significant difference between ASD and TD (*p*_*bonferonni*_ *=* 0.030, see Figure 2D) participants within Mixed Dyads; no other comparison was significant (*ps > 0*.*118*).

### EEG Hyperscanning results - Within dyads comparisons

#### Spontaneous Imitation vs Observation Phase within Control dyads

The cluster-based analysis over the 5 frequency bands of interest (Theta, Low-Alpha, High-Alpha, Beta, and Gamma) revealed a significant positive cluster in the Low-Alpha band, highlighting increased inter-brain synchrony during *Spontaneous Imitation* compared to the *Observation Phase* (*p =* 0.012, see Figure 3A) and a significant negative cluster in the High-Alpha band (*p =* 0.015, see Figure 3B) showing reduced inter-brain synchrony during *Spontaneous Imitation* compared to the *Observation Phase* within Control dyads.

**Figure 3.**
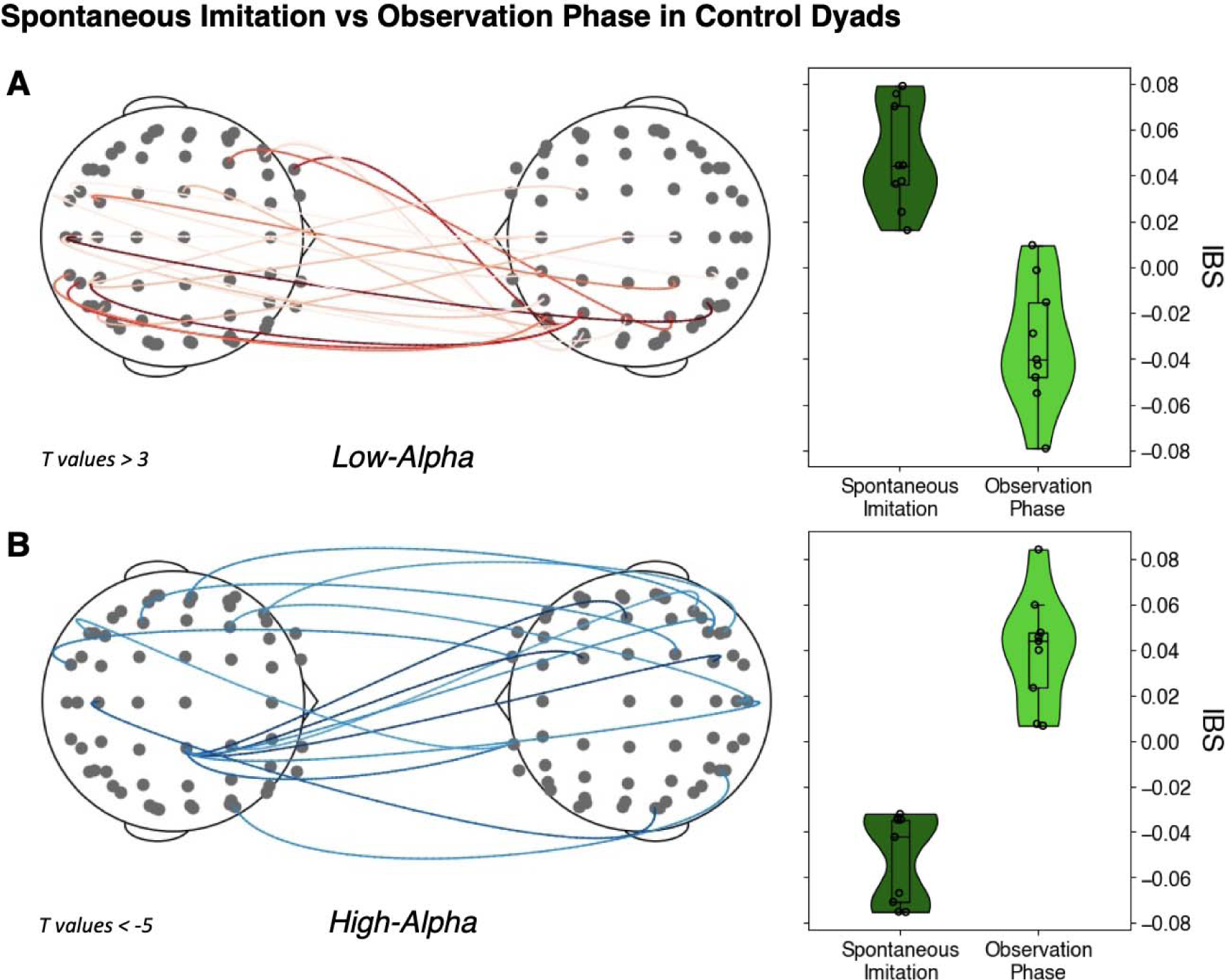
Significant inter-brain synchrony differences between *Spontaneous Imitation* and *Observation Phase* in Control Dyads revealed by cluster-based analyses in the **A**. Low-Alpha and **B**. High-Alpha frequency bands.

No other cluster-based analysis (i.e., Video Imitation vs Spontaneous Imitation within Control dyads, Spontaneous Imitation vs Observation Phase within Mixed dyads, and Video Imitation vs Spontaneous Imitation within Mixed dyads) over the 5 frequency bands of interest (Theta, Low-Alpha, High-Alpha, Beta, and Gamma) revealed any difference in inter-brain synchrony.

### EEG Hyperscanning results - Between dyads comparisons

#### Control dyads vs Mixed Dyads in the Observation Phase

No cluster-based analysis over the 5 frequency bands of interest (Theta, Low-Alpha, High-Alpha, Beta, and Gamma) revealed any difference in inter-brain synchrony between Control and Mixed Dyads in the *Observation Phase*.

#### Control Dyads vs Mixed Dyads in the Video Imitation

The cluster-based analysis over the 5 frequency bands of interest (Theta, Low-Alpha, High-Alpha, Beta, and Gamma) revealed a significant positive cluster in the Low-Alpha band, highlighting increased inter-brain synchrony for Control Dyads compared to Mixed Dyads in the *Video Imitation* condition (*p =* 0.035, see Figure 4A)

**Figure 4.**
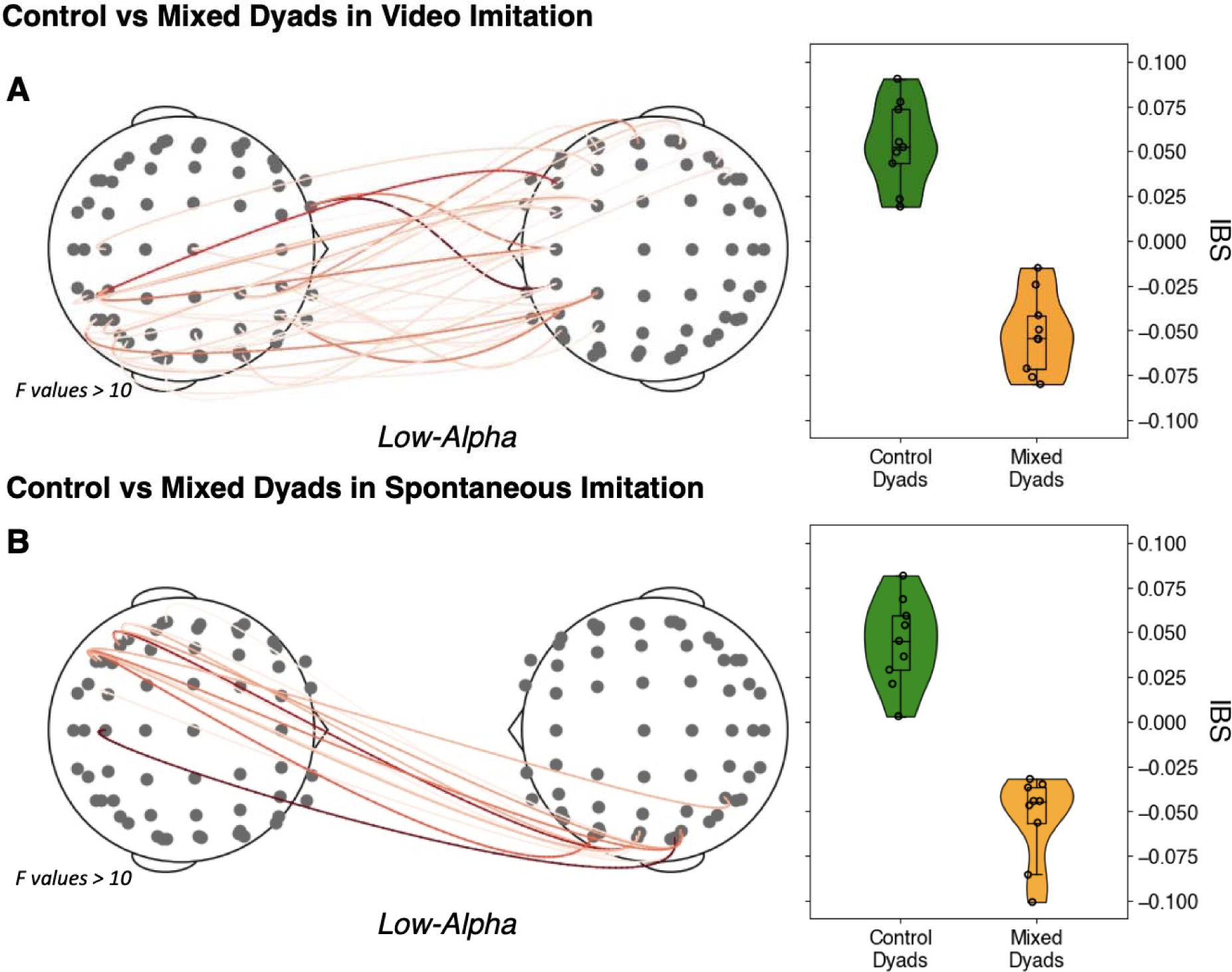
Significant inter-brain synchrony differences in the Low-Alpha band between Control and Mixed Dyads during **A**. *Video Imitation* and **B**. *Spontaneous Imitation* conditions.

#### Control dyads vs Mixed Dyads in the Spontaneous Imitation

The cluster-based analysis over the 5 frequency bands of interest (Theta, Low-Alpha, High-Alpha, Beta, and Gamma) revealed a significant positive cluster in the Low-Alpha band, highlighting increased inter-brain synchrony for Control Dyads compared to Mixed Dyads in the *Spontaneous Imitation* condition (*p =* 0.034, see Figure 4B)

### EEG Hyperscanning results - Between dyads comparisons with leader-follower distinction

The cluster-based analysis over the 5 frequency bands of interest (Theta, Low-Alpha, High-Alpha, Beta, and Gamma) revealed two significant positive clusters in the Low-Alpha and the Beta bands, highlighting increased inter-brain synchrony for Control Dyads compared to Mixed Dyads (*p =* 0.003 and *p =* 0.026, see Figure 5A and 5C), as well as a cluster in the High-Alpha band showing larger IBS for Mixed Dyads compared to Control Dyads (*p =* 0.019, see Figure 5B).

**Figure 5.**
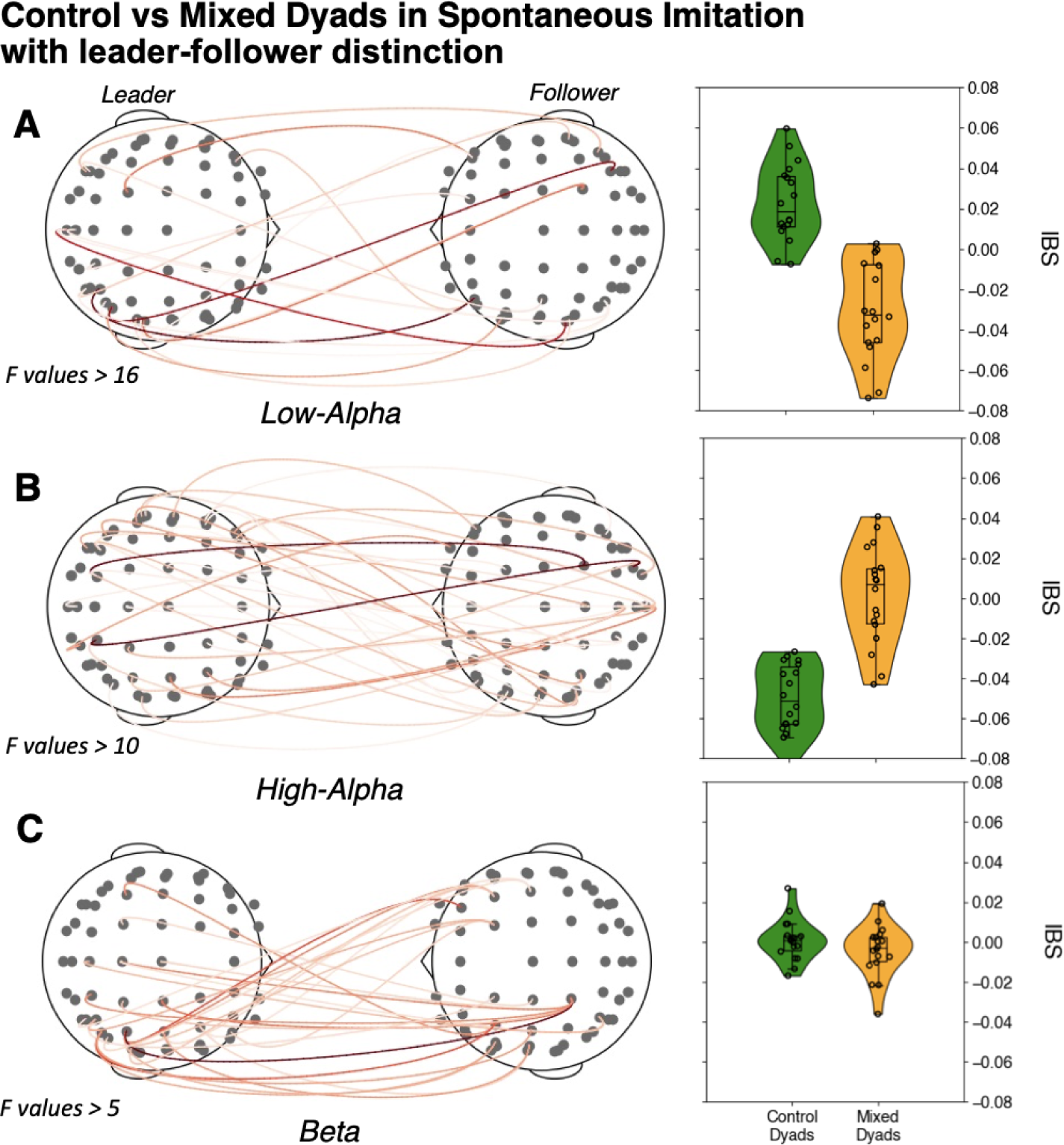
Significant inter-brain synchrony differences in the **A**. Low-Alpha; **B**. High-Alpha and **C**. Beta bands between Control and Mixed Dyads the *Spontaneous Imitation* with leader-follower distinction.

## Discussion

In this paper, we conducted dyadic social imitation experiments while recording movement kinematics and EEG-hyperscanning. Our aim was to compare the results between dyads made of two TD participants, and mixed dyads comprising ASD and TD people, with the primary hypothesis that the ASD-TD dyads would exhibit different patterns in both behavioral and inter-brain dynamics compared to TD-TD dyads.

### Behavioral differences in ecological imitating scenarios between Mixed and Control dyads

First, our research sheds new light on the behavior of individuals with ASD in dyadic social scenarios. Our findings indicate that, while there were no significant differences in overall dyadic performance between Mixed and Control dyads (i.e., *Overall Imitation* and *Synchrony*, see Figure 2A and 2B), there were notable differences in within-dyad dynamics. Specifically, individuals with ASD exhibited a preference for not taking the lead role in imitating scenarios. First, although not significant, we observe an imbalance in *Role Symmetry* (Figure 2C) that is further confirmed by breaking down the role of each individual within the dyad (Figure 2D), where we see that in Mixed dyads, participants with ASD were significantly more likely to be followers than leaders during *Spontaneous* imitation phases. This suggests that individuals with ASD may have intact lower-level skills for interpersonal imitation (Wadsworth et al., 2017), but they may be less inclined to assume leadership roles in social interactions involving TD participants. Importantly, our study underscores the importance of studying interpersonal phenomena in dynamic and ecological settings. Indeed, our findings suggest an atypical pattern at the dynamic interactive level. Previous research has often focused on pseudo-social paradigms and these scenarios may not fully capture the nuances of social interactions. By considering the ecological context in which social interactions occur, we can better understand the behavior of individuals with ASD (Schilbach et al., 2013; Dumas, 2022; Nadel, 2014b).

### Hyperscanning reveals inter-brain synchrony differences within Control and Mixed dyads

Hyperscanning provides a novel perspective on ASD, by capturing the neural activity during actual social interactions rather than in isolated contexts. Our results here show that dyads involving ASD individuals exhibit distinct patterns of brain activity compared to neurotypical dyads. First, when comparing *Spontaneous Imitation* data with the *Observation Phase*, we show a significant increase of IBS in the low-alpha band and an IBS decrease in the high-alpha band in Control dyads (Figures 3A and 3B). Although a lack of an effect does not constitute sufficient proof, we note that no such pattern has been detected in Mixed dyads. Regarding the interpretation of the results, we observe a dissociation between low- and high-alpha bands (Tognoli et al., 2007; Dumas et al., 2014). We show an increase of inter-brain synchrony during a more demanding social task in the lower band, in phase with previous results (Dumas et al., 2010; Konvalinka et al., 2014), and more IBS during the simultaneous passive viewing of videos in the high-alpha band. While an increase in high-alpha power at the intra-brain level is in line with increased visual attention accounts (Lobier et al., 2018; Peylo et al., 2021), the observation of increased IBS might be the byproduct of simultaneously processing similar stimuli (Hasson et al., 2004; Haxby et al., 2020).

### Low-Alpha IBS differentiates Control and Mixed dyads

The comparisons between Control and Mixed dyads in both *Video* and *Spontaneous* imitations show both larger inter-brain synchrony in the low-alpha band for TD-TD vs ASD-TD dyads (Figure 4A and 4B). These results are further confirmed in the comparisons with leader-follower distinction (Figure 5A), with larger low-alpha IBS for Controls compared to Mixed dyads. The leader-follower analysis also revealed patterns that were not detected in the sole condition comparison: higher IBS in the beta band for Controls (Figure 5C), in line with previous results (Dumas et al., 2010), but crucially higher IBS in the high-alpha band for Mixed dyads (Figure 5B). The different patterns of IBS, rather than a mere ‘lower’ IBS in ASD-TD dyads, suggest that distinct interactive experiences (i.e., differences in interpersonal dynamics, as suggested by our behavioral results) lead to distinguishable inter-brain neurophysiological patterns.

## Limitations

While our study provides novel insights, there are some limitations to our findings. Firstly, the sample size of our study was relatively small, with only nine dyads in each group. Therefore, caution should be exercised when generalizing our results to a larger population. Secondly, the sample of individuals with ASD in our study inherently leads to heterogeneity in terms of their clinical presentation, which could have influenced our results. Finally, while our study utilized hyperscanning to capture inter-brain dynamics during social interaction, this method has some inherent limitations such as the difficulty in establishing the causality or directionality of the observed neural patterns (Moreau and Dumas, 2021).

## Conclusion

In conclusion, our study provides new insights into the behavioral and neural differences between people with and without ASD in social imitation tasks. Our findings highlight the importance of considering the dynamic and ecological nature of social interactions, as well as the use of hyperscanning methods to capture inter-brain dynamics during actual social interactions. We also show that individuals with ASD may have intact abilities for interpersonal imitation, but still rather not take the lead in social situations involving TD participants. These differences are also highlighted at the inter-brain level, with consistent differences in inter-brain synchrony between Mixed and Control dyads in the low-alpha band. Overall, our study contributes to the growing body of literature aimed at better understanding the social cognitive processes and neural mechanisms underlying social interaction in individuals with and without ASD and provides potential markers for inter-personal approaches to psychiatric conditions with specific social misattunement (Bolis et al., 2022; Dumas, 2022).

## Acknowledgments

We thank Robert Soussignan and Emeline Mercier for their help in the manual indexing, Laurent Hugueville for his assistance in the setting of the hyperscanning system, and Florence Bouchet for her generous help in the EEG preparation.

## Fundings

G.D. was supported by the Orange Foundation for Autism Spectrum Disorders and the Fonds de recherche du Québec (FRQ; 285289). This study was supported by the Institute for Data Valorization, Montreal (IVADO; CF00137433). This study was enabled in part by support provided by Calcul Québec (www.calculquebec.ca) and Digital Research Alliance of Canada (www.alliancecan.ca). Q.M. was supported by the Unifying Neuroscience and Artificial Intelligence - Québec (UNIQUE) postdoctoral excellence fellowship.

## Notes

### Competing Interest Statement

The authors have declared no competing interest.

